## Supplementary material for "Computational investigation suggests that the cell adhesion sub-proteome is enriched for sites of pH-dependence and charge burial, whilst some key intracellular pathways may be shielded from pH fluctuations": S1 Fig

| GO-ID | GO-NAME | pkcalc-int-15-2 | pkcalc-int-nodh-3 | pkcalc-int-15-3 | pkcalc-int-5-2 | propka-range-15 | propka-range-5 | pkcalc-15-3-propka-15 | pkcalc-15-2-propka-15 | pkcalc-5-2-propka-5 | pkcalc-deltaQ-10 |
|---|---|---|---|---|---|---|---|---|---|---|---|
| GO:0021637 | trigeminal nerve structural organization | | | | | | | | | | |
| GO:0021636 | trigeminal nerve morphogenesis | | | | | | | | | | |
| GO:0007193 | adenylate cyclase-inhibiting G protein-coupled r | | | | | | | | | | |
| GO:0007186 | G protein-coupled receptor signaling pathway | | | | | | | | | | |
| GO:0098664 | G protein-coupled serotonin receptor signaling p | | | | | | | | | | |
| GO:0007196 | adenylate cyclase-inhibiting G protein-coupled g | | | | | | | | | | |
| GO:0007216 | G protein-coupled glutamate receptor signaling | | | | | | | | | | |
| GO:0007215 | glutamate receptor signaling pathway | | | | | | | | | | |
| GO:0007160 | cell-matrix adhesion | | | | | | | | | | |
| GO:0032483 | regulation of Rab protein signal transduction | | | | | | | | | | |
| GO:0051056 | regulation of small GTPase mediated signal trans | | | | | | | | | | |
| GO:0050955 | thermoception | | | | | | | | | | |
| GO:0007600 | sensory perception | | | | | | | | | | |
| GO:0050906 | detection of stimulus involved in sensory percep | | | | | | | | | | |
| GO:0051606 | detection of stimulus | | | | | | | | | | |
| GO:0032253 | dense core granule localization | | | | | | | | | | |
| GO:0032252 | secretory granule localization | | | | | | | | | | |
| GO:0007043 | cell-cell junction assembly | | | | | | | | | | |
| GO:0034329 | cell junction assembly | | | | | | | | | | |
| GO:0015793 | glycerol transmembrane transport | | | | | | | | | | |
| GO:0098609 | cell-cell adhesion | | | | | | | | | | |
| GO:0086001 | cardiac muscle cell action potential | | | | | | | | | | |
| GO:0001508 | action potential | | | | | | | | | | |
| GO:0051899 | membrane depolarization | | | | | | | | | | |
| GO:0009593 | detection of chemical stimulus | | | | | | | | | | |
| GO:0035589 | G protein-coupled purinergic nucleotide recepto | | | | | | | | | | |
| GO:0035590 | purinergic nucleotide receptor signaling pathway | | | | | | | | | | |
| GO:0050911 | detection of chemical stimulus involved in senso | | | | | | | | | | |
| GO:0050907 | detection of chemical stimulus involved in senso | | | | | | | | | | |
| GO:0007606 | sensory perception of chemical stimulus | | | | | | | | | | |
| GO:0007608 | sensory perception of smell | | | | | | | | | | |
| GO:0016339 | calcium-dependent cell-cell adhesion via plasma | | | | | | | | | | |
| GO:0098742 | cell-cell adhesion via plasma-membrane adhesic | | | | | | | | | | |
| GO:0007187 | G protein-coupled receptor signaling pathway co | | | | | | | | | | |
| GO:0099505 | regulation of presynaptic membrane potential | | | | | | | | | | |
| GO:0140627 | ubiquitin-dependent protein catabolic process v | | | | | | | | | | |
| GO:0051016 | barbed-end actin filament capping | | | | | | | | | | |
| GO:1904816 | positive regulation of protein localization to chro | | | | | | | | | | |
| GO:0097104 | postsynaptic membrane assembly | | | | | | | | | | |
| GO:0007416 | synapse assembly | | | | | | | | | | |
| GO:0007200 | phospholipase C-activating G protein-coupled re | | | | | | | | | | |
| GO:0030048 | actin filament-based movement | | | | | | | | | | |
| GO:0044331 | cell-cell adhesion mediated by cadherin | | | | | | | | | | |
| GO:0007214 | gamma-aminobutyric acid signaling pathway | | | | | | | | | | |
| GO:0045761 | regulation of adenylate cyclase activity | | | | | | | | | | |
| GO:0034332 | adherens junction organization | | | | | | | | | | |
| GO:0071880 | adenylate cyclase-activating adrenergic receptor | | | | | | | | | | |
| GO:0071875 | adrenergic receptor signaling pathway | | | | | | | | | | |
| GO:0090630 | activation of GTPase activity | | | | | | | | | | |
| GO:0001755 | neural crest cell migration | | | | | | | | | | |
| GO:0090497 | mesenchymal cell migration | | | | | | | | | | |
| GO:0014032 | neural crest cell development | | | | | | | | | | |
| GO:0050919 | negative chemotaxis | | | | | | | | | | |
| GO:0007156 | homophilic cell adhesion via plasma membrane | | | | | | | | | | |
| GO:1990806 | ligand-gated ion channel signaling pathway | | | | | | | | | | |
| GO:0071526 | semaphorin-plexin signaling pathway | | | | | | | | | | |
| GO:0032482 | Rab protein signal transduction | | | | | | | | | | |
| GO:0007190 | activation of adenylate cyclase activity | | | | | | | | | | |
| GO:0045762 | positive regulation of adenylate cyclase activity | | | | | | | | | | |
| GO:0051932 | synaptic transmission GABAergic | | | | | | | | | | |
| GO:0086002 | cardiac muscle cell action potential involved in co | | | | | | | | | | |
| GO:0086003 | cardiac muscle cell contraction | | | | | | | | | | |
| GO:0070252 | actin-mediated cell contraction | | | | | | | | | | |
| GO:1904862 | inhibitory synapse assembly | | | | | | | | | | |
| GO:0007271 | synaptic transmission cholinergic | | | | | | | | | | |
| GO:0035025 | positive regulation of Rho protein signal transdu | | | | | | | | | | |
| GO:0051057 | positive regulation of small GTPase mediated sig | | | | | | | | | | |
| GO:0035023 | regulation of Rho protein signal transduction | | | | | | | | | | |
| GO:0034472 | snRNA 3'-end processing | | | | | | | | | | |
| GO:0016180 | snRNA processing | | | | | | | | | | |
| GO:0019228 | neuronal action potential | | | | | | | | | | |
| GO:0033627 | cell adhesion mediated by integrin | | | | | | | | | | |
| GO:0098703 | calcium ion import across plasma membrane | | | | | | | | | | |
| GO:0070509 | calcium ion import | | | | | | | | | | |
| GO:1902656 | calcium ion import into cytosol | | | | | | | | | | |
| GO:0006833 | water transport | | | | | | | | | | |
| GO:0007229 | integrin-mediated signaling pathway | | | | | | | | | | |
