## Supplementary material for "Computational investigation suggests that the cell adhesion sub-proteome is enriched for sites of pH-dependence and charge burial, whilst some key intracellular pathways may be shielded from pH fluctuations": S2 Fig

| GO-ID | GO-NAME | pkcalc-int-15-2 | pkcalc-int-nodh-3 | pkcalc-int-15-3 | pkcalc-int-5-2 | propka-range-15 | propka-range-5 | pkcalc-15-3-propka-15 | pkcalc-15-2-propka-15 | pkcalc-5-2-propka-5 | pkcalc-deltaQ-10 |
|---|---|---|---|---|---|---|---|---|---|---|---|
| GO:0004888 | transmembrane signaling receptor activity | | | | | | | | | | |
| GO:0038023 | signaling receptor activity | | | | | | | | | | |
| GO:0060089 | molecular transducer activity | | | | | | | | | | |
| GO:0016494 | C-X-C chemokine receptor activity | | | | | | | | | | |
| GO:0004950 | chemokine receptor activity | | | | | | | | | | |
| GO:0001637 | G protein-coupled chemoattractant receptor act | | | | | | | | | | |
| GO:0004930 | G protein-coupled receptor activity | | | | | | | | | | |
| GO:0016493 | C-C chemokine receptor activity | | | | | | | | | | |
| GO:0016502 | nucleotide receptor activity | | | | | | | | | | |
| GO:0022829 | wide pore channel activity | | | | | | | | | | |
| GO:0005243 | gap junction channel activity | | | | | | | | | | |
| GO:0001640 | adenylate cyclase inhibiting G protein-coupled gl | | | | | | | | | | |
| GO:0098988 | G protein-coupled glutamate receptor activity | | | | | | | | | | |
| GO:0008066 | glutamate receptor activity | | | | | | | | | | |
| GO:0004983 | neuropeptide Y receptor activity | | | | | | | | | | |
| GO:0008188 | neuropeptide receptor activity | | | | | | | | | | |
| GO:0008528 | G protein-coupled peptide receptor activity | | | | | | | | | | |
| GO:0001653 | peptide receptor activity | | | | | | | | | | |
| GO:0005267 | potassium channel activity | | | | | | | | | | |
| GO:0005227 | calcium-activated cation channel activity | | | | | | | | | | |
| GO:0022839 | monoatomic ion-gated channel activity | | | | | | | | | | |
| GO:0005245 | voltage-gated calcium channel activity | | | | | | | | | | |
| GO:0022843 | voltage-gated monoatomic cation channel activi | | | | | | | | | | |
| GO:0005244 | voltage-gated monoatomic ion channel activity | | | | | | | | | | |
| GO:0022832 | voltage-gated channel activity | | | | | | | | | | |
| GO:0045125 | bioactive lipid receptor activity | | | | | | | | | | |
| GO:0001594 | trace-amine receptor activity | | | | | | | | | | |
| GO:0008227 | G protein-coupled amine receptor activity | | | | | | | | | | |
| GO:0022851 | GABA-gated chloride ion channel activity | | | | | | | | | | |
| GO:0022824 | transmitter-gated monoatomic ion channel activ | | | | | | | | | | |
| GO:0005230 | extracellular ligand-gated monoatomic ion chan | | | | | | | | | | |
| GO:0022835 | transmitter-gated channel activity | | | | | | | | | | |
| GO:0030594 | neurotransmitter receptor activity | | | | | | | | | | |
| GO:0004970 | glutamate-gated receptor activity | | | | | | | | | | |
| GO:0099604 | ligand-gated calcium channel activity | | | | | | | | | | |
| GO:0022848 | acetylcholine-gated monoatomic cation-selective | | | | | | | | | | |
| GO:1904315 | transmitter-gated monoatomic ion channel activ | | | | | | | | | | |
| GO:0099529 | neurotransmitter receptor activity involved in re | | | | | | | | | | |
| GO:0098960 | postsynaptic neurotransmitter receptor activity | | | | | | | | | | |
| GO:0005231 | excitatory extracellular ligand-gated monoatomi | | | | | | | | | | |
| GO:0008569 | minus-end-directed microtubule motor activity | | | | | | | | | | |
| GO:0003777 | microtubule motor activity | | | | | | | | | | |
| GO:0003774 | cytoskeletal motor activity | | | | | | | | | | |
| GO:0033038 | bitter taste receptor activity | | | | | | | | | | |
| GO:0008527 | taste receptor activity | | | | | | | | | | |
| GO:0019956 | chemokine binding | | | | | | | | | | |
| GO:0008331 | high voltage-gated calcium channel activity | | | | | | | | | | |
| GO:0070679 | inositol 145 trisphosphate binding | | | | | | | | | | |
| GO:0005549 | odorant binding | | | | | | | | | | |
| GO:1901338 | catecholamine binding | | | | | | | | | | |
| GO:0005272 | sodium channel activity | | | | | | | | | | |
| GO:0015254 | glycerol channel activity | | | | | | | | | | |
| GO:0015168 | glycerol transmembrane transporter activity | | | | | | | | | | |
| GO:0016500 | protein-hormone receptor activity | | | | | | | | | | |
| GO:0004993 | G protein-coupled serotonin receptor activity | | | | | | | | | | |
| GO:0004953 | icosanoid receptor activity | | | | | | | | | | |
| GO:0016907 | G protein-coupled acetylcholine receptor activity | | | | | | | | | | |
| GO:0099528 | G protein-coupled neurotransmitter receptor ac | | | | | | | | | | |
| GO:0015464 | acetylcholine receptor activity | | | | | | | | | | |
| GO:0004935 | adrenergic receptor activity | | | | | | | | | | |
| GO:0004936 | alpha-adrenergic receptor activity | | | | | | | | | | |
| GO:0098639 | collagen binding involved in cell-matrix adhesior | | | | | | | | | | |
| GO:0005249 | voltage-gated potassium channel activity | | | | | | | | | | |
| GO:0005223 | intracellularly cGMP-activated cation channel ac | | | | | | | | | | |
| GO:0005221 | intracellularly cyclic nucleotide-activated monoa | | | | | | | | | | |
| GO:0043855 | cyclic nucleotide-activated monoatomic ion char | | | | | | | | | | |
| GO:0005217 | intracellularly ligand-gated monoatomic ion char | | | | | | | | | | |
| GO:0005222 | intracellularly cAMP-activated cation channel act | | | | | | | | | | |
| GO:0045028 | G protein-coupled purinergic nucleotide recepto | | | | | | | | | | |
| GO:0001614 | purinergic nucleotide receptor activity | | | | | | | | | | |
| GO:0004890 | GABA-A receptor activity | | | | | | | | | | |
| GO:0016917 | GABA receptor activity | | | | | | | | | | |
| GO:0004984 | olfactory receptor activity | | | | | | | | | | |
| GO:0019957 | C-C chemokine binding | | | | | | | | | | |
| GO:0030215 | semaphorin receptor binding | | | | | | | | | | |
| GO:0008574 | plus-end-directed microtubule motor activity | | | | | | | | | | |
| GO:0042923 | neuropeptide binding | | | | | | | | | | |
| GO:0015250 | water channel activity | | | | | | | | | | |
| GO:0005372 | water transmembrane transporter activity | | | | | | | | | | |
| GO:0008020 | G protein-coupled photoreceptor activity | | | | | | | | | | |
| GO:0009881 | photoreceptor activity | | | | | | | | | | |
| GO:0017154 | semaphorin receptor activity | | | | | | | | | | |
| GO:0008503 | benzodiazepine receptor activity | | | | | | | | | | |
| GO:0051378 | serotonin binding | | | | | | | | | | |
| GO:0061608 | nuclear import signal receptor activity | | | | | | | | | | |
| GO:0140142 | nucleocytoplasmic carrier activity | | | | | | | | | | |
| GO:0004955 | prostaglandin receptor activity | | | | | | | | | | |
| GO:0004954 | prostanoid receptor activity | | | | | | | | | | |
| GO:0000146 | microfilament motor activity | | | | | | | | | | |
| GO:0005544 | calcium-dependent phospholipid binding | | | | | | | | | | |
| GO:0005545 | 1-phosphatidylinositol binding | | | | | | | | | | |
| GO:0005085 | guanyl-nucleotide exchange factor activity | | | | | | | | | | |
| GO:0030695 | GTPase regulator activity | | | | | | | | | | |
| GO:0060589 | nucleoside-triphosphatase regulator activity | | | | | | | | | | |
| GO:0005096 | GTPase activator activity | | | | | | | | | | |
| GO:0008381 | mechanosensitive monoatomic ion channel activ | | | | | | | | | | |
| GO:0005251 | delayed rectifier potassium channel activity | | | | | | | | | | |
| GO:0005041 | low-density lipoprotein particle receptor activity | | | | | | | | | | |
| GO:0045294 | alpha-catenin binding | | | | | | | | | | |
| GO:0098973 | structural constituent of postsynaptic actin cytos | | | | | | | | | | |
| GO:0003707 | nuclear steroid receptor activity | | | | | | | | | | |
| GO:0005248 | voltage-gated sodium channel activity | | | | | | | | | | |
| GO:1990756 | ubiquitin-like ligase-substrate adaptor activity | | | | | | | | | | |
| GO:0140767 | enzyme-substrate adaptor activity | | | | | | | | | | |
| GO:0140662 | ATP-dependent protein folding chaperone | | | | | | | | | | |
| GO:0051015 | actin filament binding | | | | | | | | | | |
| GO:0045499 | chemorepellent activity | | | | | | | | | | |
| GO:0030020 | extracellular matrix structural constituent confer | | | | | | | | | | |
| GO:0005547 | phosphatidylinositol-345-trisphosphate binding | | | | | | | | | | |
| GO:0008013 | beta-catenin binding | | | | | | | | | | |
| GO:0032934 | sterol binding | | | | | | | | | | |
